# JSTA: joint cell segmentation and cell type annotation for spatial transcriptomics

**DOI:** 10.1101/2020.09.18.304147

**Authors:** Russell Littman, Zachary Hemminger, Robert Foreman, Douglas Arneson, Guanglin Zhang, Fernando Gómez-Pinilla, Xia Yang, Roy Wollman

## Abstract

RNA hybridization based spatial transcriptomics provides unparalleled detection sensitivity. However, inaccuracies in segmentation of image volumes into cells cause misassignment of mRNAs which is a major source of errors. Here we develop JSTA, a computational framework for Joint cell Segmentation and cell Type Annotation that utilizes prior knowledge of cell-type specific gene expression. Simulation results show that leveraging existing cell type taxonomy increases RNA assignment accuracy by more than 45%. Using JSTA we were able to classify cells in the mouse hippocampus into 133 (sub)types revealing the spatial organization of CA1, CA3, and Sst neuron subtypes. Analysis of within cell subtype spatial differential gene expression of 80 candidate genes identified 43 with statistically significant spatial differential gene expression across 61 (sub)types. Overall, our work demonstrates that known cell type expression patterns can be leveraged to improve the accuracy of RNA hybridization based spatial transcriptomics while providing highly granular cell (sub)type information. The large number of newly discovered spatial gene expression patterns substantiates the need for accurate spatial transcriptomics measurements that can provide information beyond cell (sub)type labels.

## Introduction

Spatial transcriptomics has been employed to explore the spatial and cell-type specific gene expression to better understand physiology and disease^1–7^. Compared to other spatial transcriptomics methods, RNA hybridization based approaches provided the highest RNA detection accuracies with capture rates > 95%^8^. With the development of combinatorial approaches for RNA hybridization, the ability to measure the expression of hundreds to thousands of genes makes hybridization based methods an attractive platform for spatial transcriptomics^8–13^. Nonetheless, unlike dissociative approaches, such as single-cell RNA sequencing (scRNAseq) where cells are captured individually, RNA hybridization based approaches have no a priori information of which cell a measured RNA molecule belongs to. Segmentation of image volumes into cells is therefore required to convert RNA detection into spatial single-cell data. Assigning mRNA to cells remains a challenging problem that can substantially compromise the overall accuracy of combinatorial FISH approaches.

Generation of spatial single-cell data from imaging based spatial transcriptomics relies on algorithmic segmentation of images into cells. Current combinatorial FISH work uses watershed based algorithms with nuclei as seeds, and the total mRNA density to establish cell borders^12,13,14^. Watershed algorithm was proposed more than 40 years ago^15^ and newer segmentation algorithms that utilize state of the art machine learning approaches have been shown to improve upon classical watershed approach ^16–20^. However, their performance is inherently bounded by the quality of the “ground truth” dataset used for training. In tissue regions with dense cell distributions, there is simply not enough information in the images to perform accurate manual labeling and create a sufficiently accurate ground truth training datasets. Therefore, there is an urgent need for new approaches that can combine image information with external datasets to improve image segmentation and thereby the overall accuracy of spatial transcriptomics.

Due to the deficiency in existing image segmentation algorithms, a few segmentation free spatial transcriptomics approaches were proposed. pciSeq assigns cell types to nuclei based on proximity to mRNA of marker genes, circumventing the need for pixel level segmentation^9^. Similarly, SSAM creates cell type maps based on RNA distributions ignoring cellular boundaries^21^. However, both pciSeq and SSAM are limited to cell type information and do not create a segmentation map for the assignment of non-cell-type marker genes. Therefore while both pciSeq and SSAM leverage cell type catalogs to provide insights into the spatial distribution of different cell types they do not produce a high quality cell segmentation map, are limited to cell (sub)type label information, and fail to assign all mRNAs to cells.

Here we present JSTA, a computational framework for jointly determining cell (sub)types and assigning mRNAs to cells by leveraging previously defined cell types through scRNAseq. Our approach relies on maximizing the internal consistency of pixel assignment into cells to match known expression patterns. We compared JSTA to watershed in assigning mRNAs to cells through simulation studies to evaluate their accuracy. Application of JSTA to MERFISH measurements of gene expression in the mouse hippocampus together with Neocortical Cell Type Taxonomy^22^ (NCTT) provides a highly granular map of cell (sub)type spatial organization and identified many spatially differentially expressed genes (spDEGs) within these (sub)types^23^.

## Results

### JSTA overview and method

Our computational framework of JSTA is based on improving initial watershed segmentation by incorporating cell (sub)type probabilities for each pixel and iteratively adjusting the assignment of boundary pixels based on those probabilities (Figure 1a).

**Figure 1.**
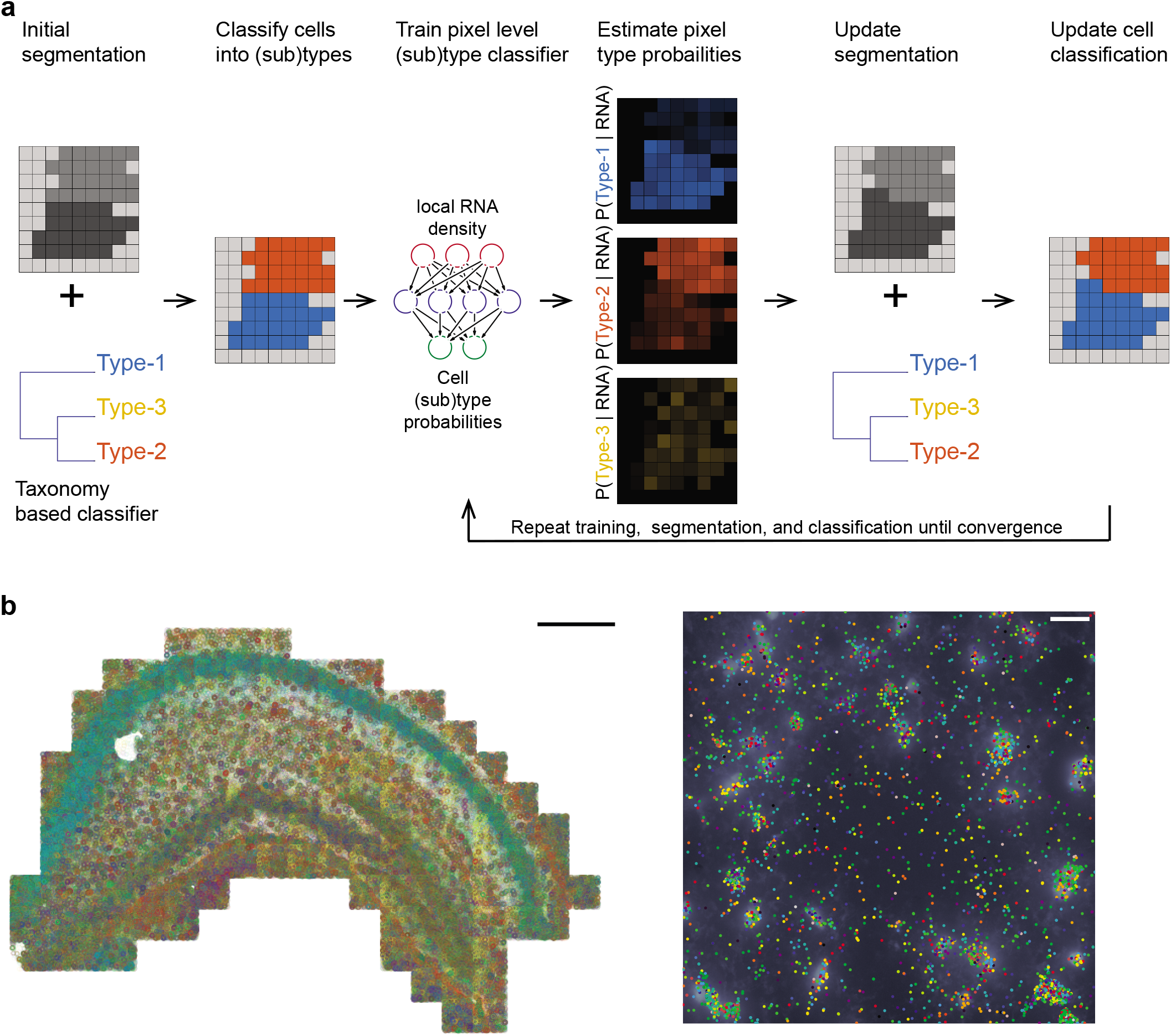
Overview of JSTA and the spatial transcriptomics data used for performance evaluation. **a**. JSTA overview. Initially, watershed based segmentation is performed and a cell level type classifier is trained based on the NCTT data. The cell level classifier then assigns cell (sub)types (red and blue in this cartoon example). Based on the current assignment of pixels to cell (sub)types, a new DNN is trained to estimate the probabilities that each pixel comes from each of the possible (sub)types given the local RNA density at each pixel. In this example, two pixels that were initially assigned to the “red” cells got higher probability to be of a blue type. Since the neighbor cell is of type “blue” they were reassigned to that cell during segmentation update. Using the updated segmentation and the cell type classifier cell types are reassigned. The tasks of training, segmentation, and classification are repeated over many iterations until convergence. **b**. Multiplexed Error Robust Fluorescent in situ hybridization (MERFISH) and DAPI stained nuclei in the mouse hippocampus. Each gene is represented by a different color. For the entire hippocampus (left), only the mRNA spots are shown with a scale bar of 500 microns. On the zoomed-in section (right), each gene is represented by a different color dot, and the DAPI intensity is displayed in white. The scale bar is 20 microns.

To evaluate JSTA we chose to use the mouse hippocampus for two reasons. 1) the mouse hippocampus has high cell (sub)type diversity as it includes more than 35% of all cell (sub)types defined by the NCTT. 2) the mouse hippocampus has areas of high and low cell density. These two reasons make the mouse hippocampus a good test case for the hypothesis that external cell (sub)type specific expression data could be leveraged to increase the accuracy of spatial transcriptomics, as implemented in our approach. We performed Multiplexed Error Robust Fluorescent In Situ Hybridization (MERFISH) of 163 genes which include 83 marker genes and 80 non-marker genes previously implicated with biological importance in traumatic brain injury (Figure 1b). Combining this MERFISH dataset, DAPI stained nuclei, and the NCTT reference dataset using JSTA, we created a segmentation map that assigns all mRNAs to cells while simultaneously classifying all cells into granular (sub)types based on NCTT.

In JSTA, we leverage the NCTT information to infer probabilities at the pixel level. However, learning these probabilities from NCTT is challenging for two reasons. 1) NCTT data was acquired with scRNAseq technology that has higher sparsity due to low capture rates and needs to be harmonized. 2) NCTT data provides expression patterns at the cell level and not the pixel level. We expect the mean expression among all pixels in a cell to be the same as that of the whole cell. Yet, variance and potentially higher distribution moments of the pixel level distribution are likely different from those of the cell level distribution due to sampling and biological factors such as variability in subcellular localization of mRNA molecules^13^. To address these issues JSTA learns the pixel level cell (sub)type probabilities using two distinct deep neural networks (DNN) classifiers, a cell level type classifier, and a pixel type classifier. Overall, JSTA learns three distinct layers of information: segmentation map, pixel level classifier, and cell level classifier.

Learning of model parameters is done using a combination of NCTT and the MERFISH data. The cell type classifier is learned directly from NCTT data after harmonization. The other two layers are learned iteratively using expectation maximization (EM) approach^24^. Given the current cell type assignment to cells, we train a pixel level DNN classifier to output the cell (sub)type probability of each pixel. The inputs for the pixel level classifier are the local mRNA density of all marker genes at these pixels. The updated pixel classifier is used to assign probabilities to all border pixels. The new probabilities are then used to “flip” border pixels’ assignment based on their type probabilities. The updating of the segmentation map requires an update of the cell level type classification which triggers a need for an update of pixel level classifier training. This process is then repeated until convergence. Analysis of the mean pixel level cell (sub)type classification accuracy shows an increase in the algorithm’s classification confidence over time demonstrating that the NCTT external information gets iteratively incorporated into the tasks of cell segmentation and type annotation (Fig S1). For computational efficiency, we iterate between training, reassignment, and reclassification in variable rates. As this approach uses cell type information to improve border assignment between neighboring cells, in cases where two neighboring cells are of the same type, the border between them will stay the same as the initial watershed segmentation. The final result is a cell type segmentation map that is initialized based on watershed and adjusted to allow pixels to be assigned to cells to maximize consistency between local RNA density and cell type expression priors.

### Performance evaluations

#### Performance evaluation using simulated hippocampus data

To test the performance of our approach we utilized synthetic data generated based on the NCTT^23^ (Figure 2ab). Details on the synthetic generation of cell position, morphologies, type, and expression profiles are available in the methods section. Using this synthetic data we evaluated the performance of JSTA in comparison to watershed at different cell type granularities. For example, two cells next to each other that are of subtypes CA1sp1 and CA1sp4 would add to the error in segmentation, but if the cell type resolution decreases to CA1 cells, these would be considered the same type, and misassignment of mRNA between these cells is no longer penalized. Evaluating the methods in this manner allows us to explore the trade-off between cell type granularity and mRNA assignment accuracy. Our analysis shows that JSTA consistently outperforms watershed at assigning spots to cells (Figure 2c). Interestingly, the benefit of JSTA was evident even with a small number of genes (Figure 2d). With just 12 genes, the performance jumps to 0.50 at the highest cell type granularity, which is already higher than watershed’s accuracy; at a granularity of 16 cell types, the accuracy reached 0.62 (Figure 2cd). Overall the synthetic data showed that JSTA outperforms watershed approach, and at physiologically relevant parameters, can increase mRNA assignment accuracy by > 45%.

**Figure 2.**
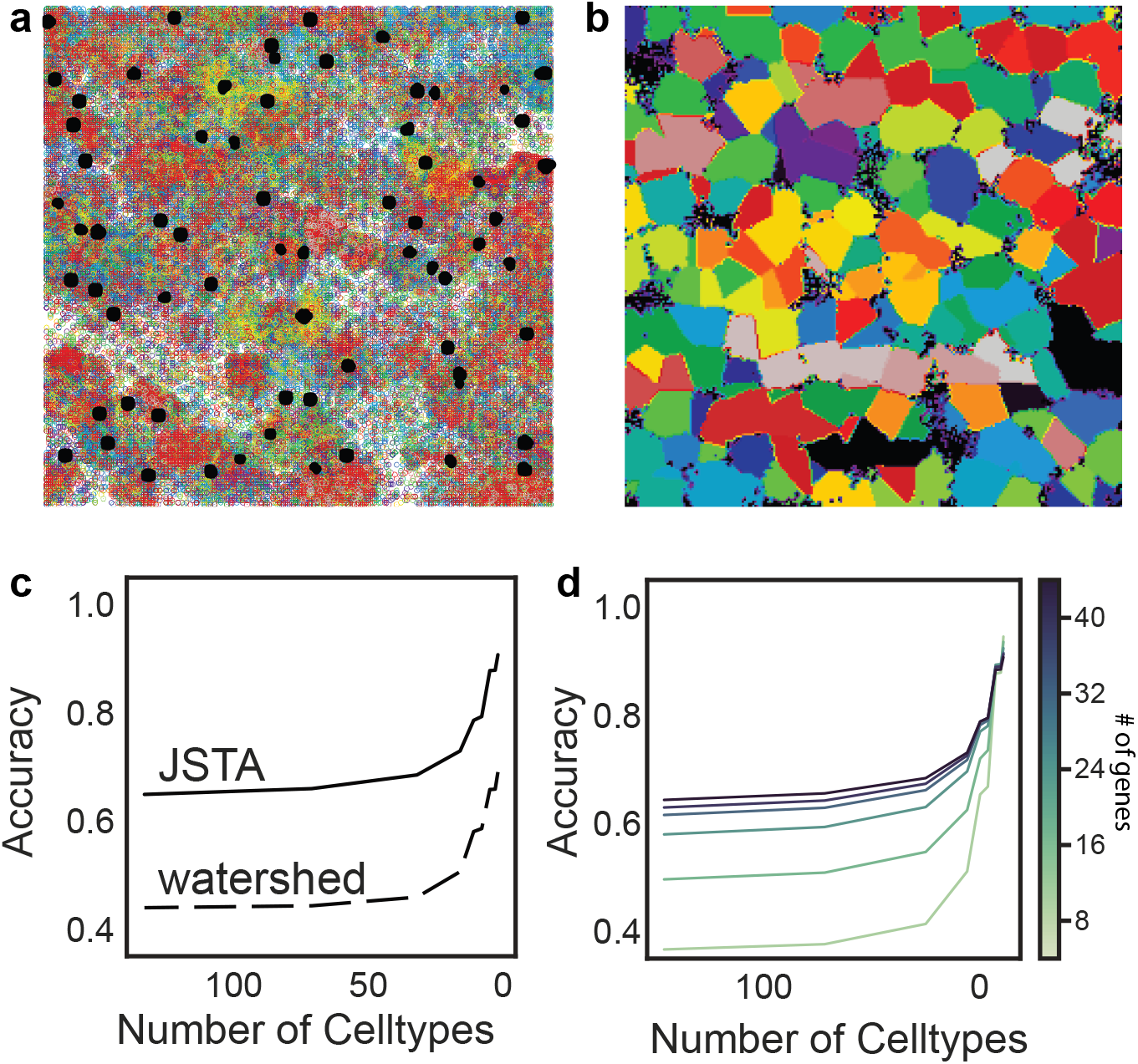
Performance evaluation of JSTA using simulated data. **a**. Representative synthetic dataset of nuclei (black) and mRNAs, where each color represents a different gene. **b**. Ground truth boundaries of the cells. Each color represents a different cell. **c**. Average Accuracy of calling mRNA spots to cells at different cell type resolutions using 83 genes. Accuracy was determined by the assignment of each mRNA molecule to the correct cell type. JSTA (solid line) is more accurate than Watershed (dashed line) at assigning mRNA molecules to the correct cells (FDR < 0.05). Statistical significance was determined with a Mann-Whitney test and false discovery rate correction. **d**. Accuracy of calling mRNA spots to cells when using JSTA to segment cells with a lower number of genes (8-44 genes tested). The color of the line gets progressively darker as the number of genes used increases.

#### Performance evaluation using empirical spatial transcriptomics of mouse hippocampus

We next tested the performance of JSTA using empirical data and evaluated its ability to recover the known spatial distribution of coarse neuron types across the hippocampus (Figure 3). First, we subset the NCTT scRNAseq data to the shared cell type marker genes we have in our MERFISH data and harmonized the MERFISH and scRNAseq datasets^25^. Using the cell type annotations from the single-cell data, we trained a DNN to classify cell types. As expected, our classifier derived a cell type mapping agreeing with known spatial patterns in the hippocampus (Figure 3a). For example, CA1, CA3, and DG cells were found with high specificity to their known subregions (Figure 3b). We found that the gene expression of the segmented cells in MERFISH data highly correlated with their scRNAseq counterparts, and displayed similar correlation patterns between different cell types (Figure 3c) as seen in scRNAseq data (Figure 3d). These results show that our data and JSTA algorithm can recover existing knowledge on the spatial distribution of cell types and their gene expression patterns in the mouse hippocampus.

**Figure 3.**
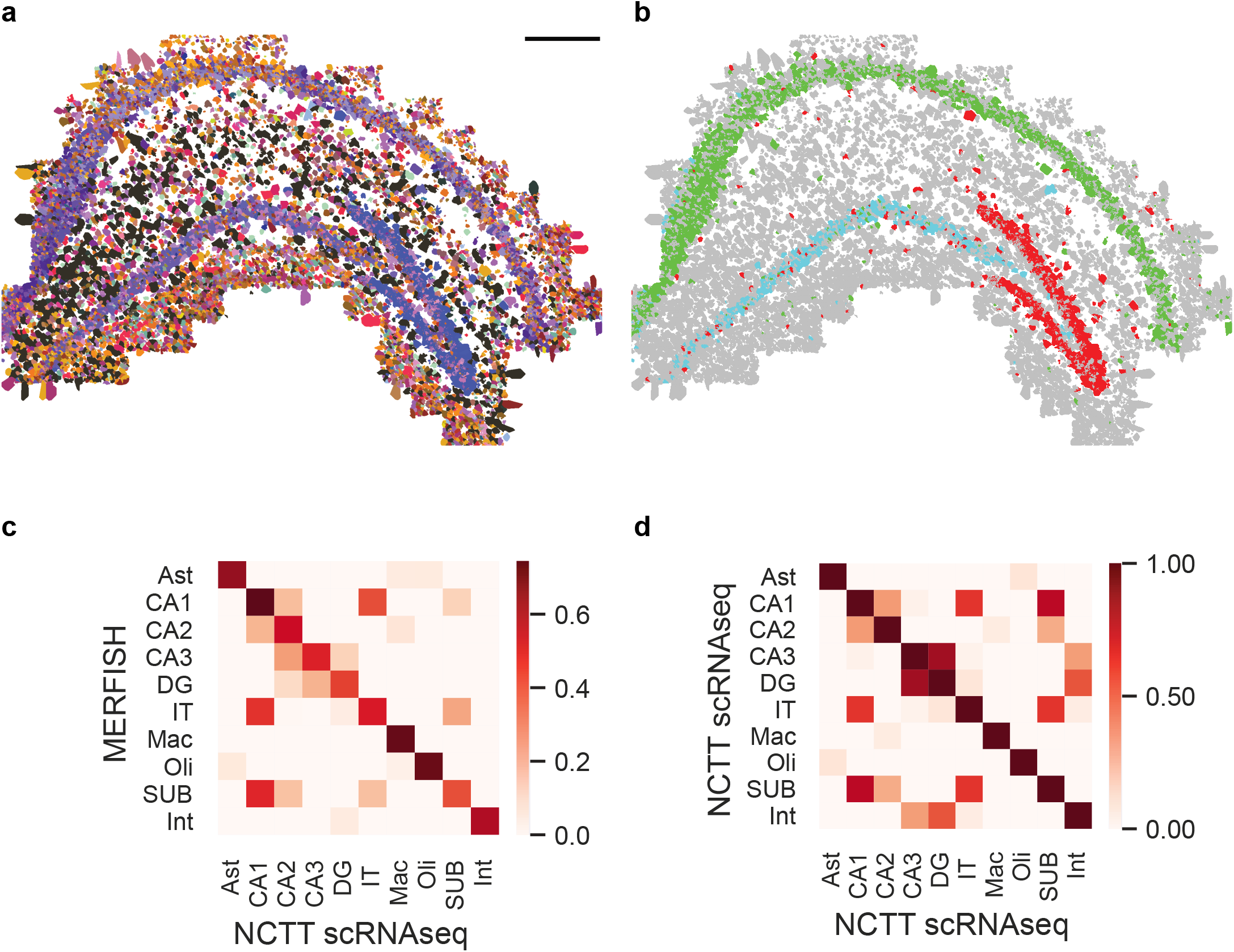
Segmentation of MERFISH data from the hippocampus using JSTA. **a**. High resolution cell type map of 133 cell (sub)types. Colors match those defined by NCTT. Scale bar is 500 microns. **b**. JSTA based classification of CA1 (green), CA3 (cyan), and DG (red) neurons matches their known domains. **c**. Correlation of 163 genes across major cell types between MERFISH measurements to scRNAseq data from NCTT. **d**. Correlation of same genes as in c between expression of types in scRNAseq data from NCTT.

### Applications of JSTA for biological discovery

#### JSTA identifies spatial distribution of highly granular cell (sub)types in the hippocampus

A key benefit of JSTA is its ability to jointly segment cells in images and classify them into highly granular cell (sub)types. Our analysis of mouse hippocampus MERFISH data found that these subtypes, defined only based on their gene expression patterns, have high spatial localization in the hippocampus. From lateral to medial hippocampus, the subtypes transitioned spatially from CA1sp10 to CA1sp6 (Figure 4a). Likewise, JSTA revealed a non-uniform distribution of subtypes in the CA3 region. From lateral to medial hippocampus, the subtypes transitioned from CA3sp4 to CA3sp6 (Figure 4b). This gradient of subtypes reveals a high level of spatial organization and points to potentially differential roles for these subtypes.

**Figure 4.**
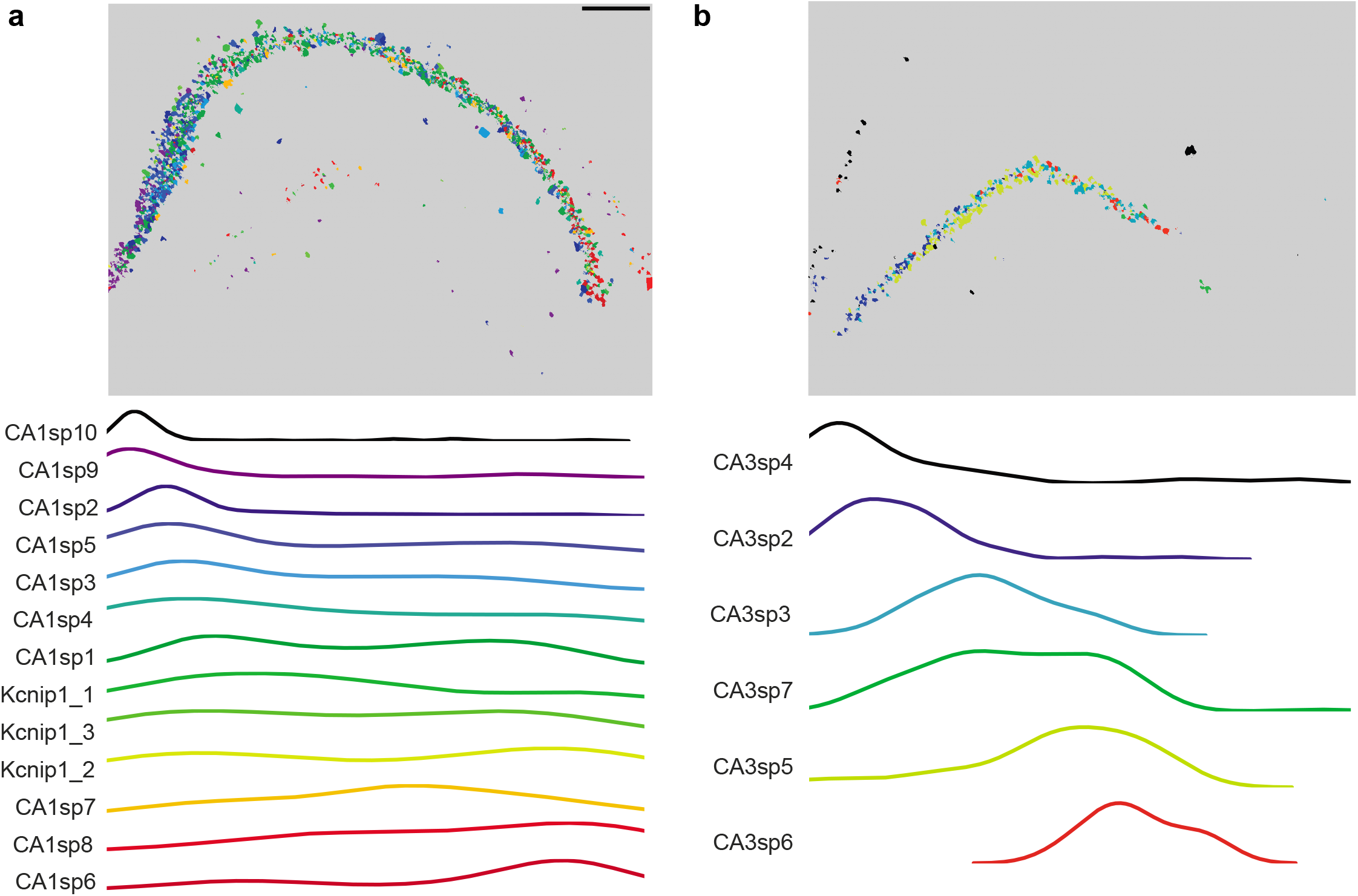
Spatial distribution of neuronal subtypes in the hippocampus. **a**. Cell subtype map of CA1 neurons in the hippocampus. Scale bar is 500 micron. Distribution of CA1 subtypes in the hippocampus, computed by projecting cell centers to the x axis. **b**. Cell subtype map of CA3 neurons in the hippocampus. Distribution of CA3 subtypes in the hippocampus, computed by projecting the cell centers to the lateral to medial axis.

#### JSTA shows that spatially proximal cell subtypes are transcriptionally similar

Next, we tested whether across different cell types spatial patterns match their expression patterns by evaluating the colocalization of cell subtypes and their transcriptional similarity. Indeed, spatially proximal CA1 subtypes showed high transcriptional similarity (Figure 5a, S2). For example, cells in the subtypes CA1sp3, CA1sp1, and CA1sp6 are proximal to each other and show a high transcriptional correlation. Interestingly, this relationship was not bidirectional, and transcriptional similarity by itself is not necessarily predictive of spatial proximity. For example, subtypes CA1sp10, CA1sp7, and CA1sp4, show >0.95 correlation but are not proximal to each other. Similar findings were seen in the CA3 region as well (Figure 5b, S2).

**Figure 5.**
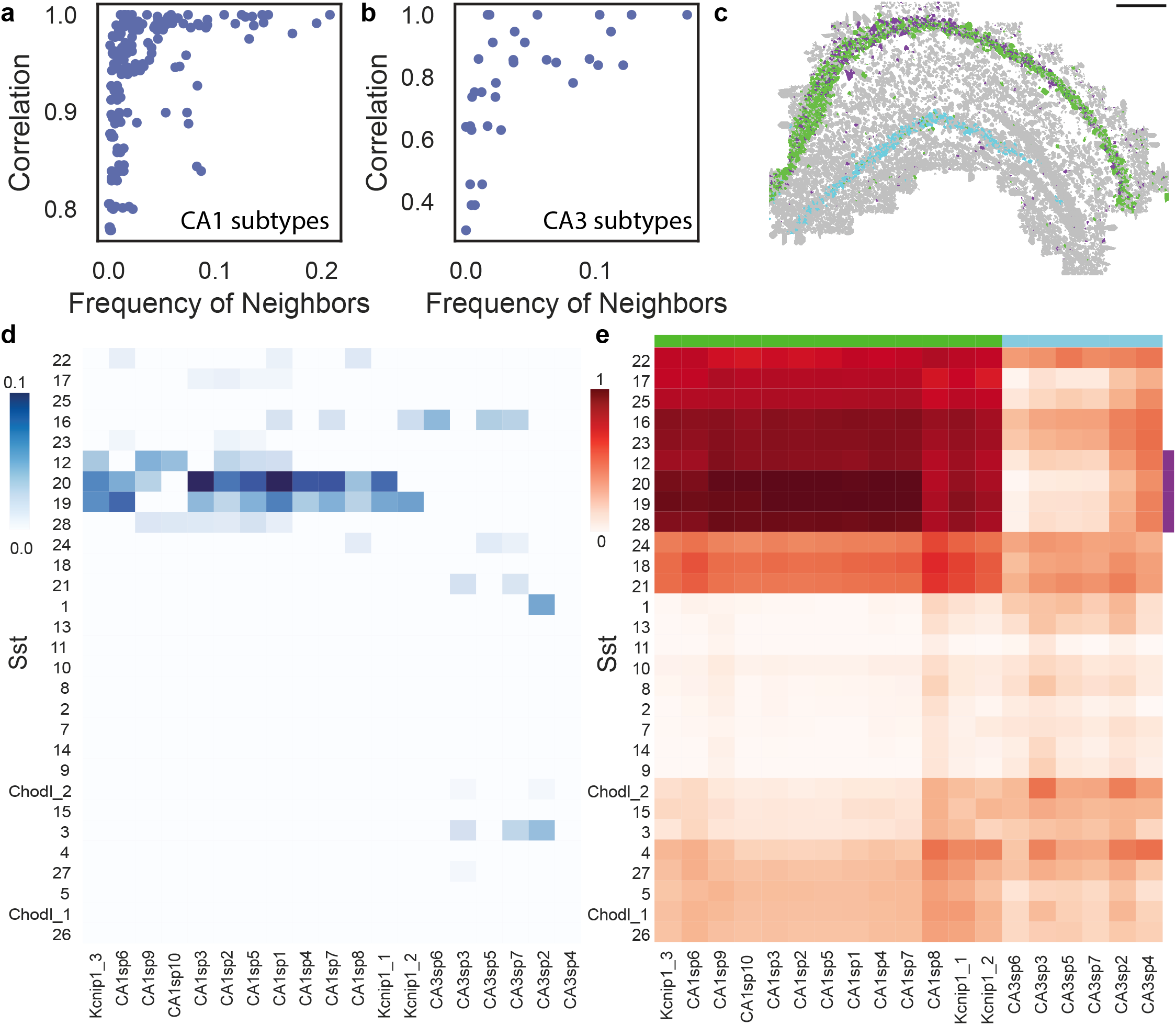
Agreement between spatial proximity and gene coexpression in high granular cell subtypes in the Hippocampus. **a-b**. Relationship between the frequency of a (sub)type’s neighbors and it’s transcriptional pearson correlation in CA1 subtypes (**a**) and CA3 subtypes (**b**). **c**. Cell type map in the hippocampus shows specific colocalization patterns between subset of Sst subtypes (purple) and CA1 neurons (green); these Sst subtypes do not colocalize with CA3 neurons (cyan). **d**. Colocalization patterns of Sst subtypes with CA1 and CA3 subtypes. Sst subtypes that colocalize with the CA1 subtypes have high transcriptional similarity. Colocalization was defined as the percent of neighbors that are of that subtype (see methods) **e** Transcriptional correlation patterns between Sst subtypes and CA1 and CA3 neurons. Green, purple and cyan sidebars highlight the subset of Sst co-localized with CA1 (purple), CA1 (green) and CA3 (cyan).

To test if this principle goes beyond subtypes of the same type we compared CA1 neurons and the Sst interneurons. We found that many Sst subtypes have high specificity in their localization and are transcriptionally related to their non Sst neighbors. Using permutation tests we found that subtypes Sst12, Sst19, Sst20, Sst28 subtypes are significantly colocalized with these same subtypes and are specific to the CA1 region (Figure 5cd, methods). Analysis of their transcriptional similarity showed that these subtypes are highly correlated in their gene expression to all CA1 subtypes (Figure 5e) but not to CA3 subtypes. These results show that both within a cell type and across cell types spatial proximity indicates similarity in expression patterns.

#### JSTA identifies spatial differential gene expression (spDEGs)

Given our results on the relationship between spatial localization and gene expression patterns across cell subtypes, we next tested whether spDEGs within the highly granular cell subtypes can be identified. We focused our analysis on the 80 genes in our dataset that were not cell marker genes used to classify cells into cell (sub)types. We identified spDEGs by determining if the spatial expression pattern of a given gene was statistically different from a null distribution by permuting the gene expression values. Importantly, the null model was restricted to the permutation of only the cells within that subtype. As a result our spDEGs analysis specifically identifies genes whose expression within a specific subtype has a spatial distribution that is different than random. We found that within hippocampal cell subtypes, many genes were differentially expressed based on their location (Figure 6). For example, *Tox* in CA1sp1 shows higher expression on the medial side of the hippocampus and decreases to the lateral side. *Leng8* in subtype CA3sp3 is highly expressed closer to the CA1 region, and lower in the medial CA3. *Hecw1* in the DG2 subtype has varying spatial distribution in the DG region. The lower portion of the DG has clusters of higher expression, while the upper portion has lower expression. These spatial differences in gene expression are not limited to neuronal subtypes. Astrocytes subtype “Astro1” shows spatial heterogeneity in expression of *Thra*, with large patches of high expression levels and other patches of little to no expression (Figure 6a). Overall, we tested for spDEGs in 61 (sub)types with more than 40 cells. We found that all 61 of the tested hippocampal cell subtypes have spDEGs (Figure 6b, S3b), with more than 50% (43 of 80) of the tested genes showing non-random spatial pattern (Figure 6c, S3c). Certain genes also show spatial patterns in many subtypes (e.g., *Thra* S3ac), while others are more specific to a one or a few subtypes (e.g., *Farp1*, S3ac).

**Figure 6.**
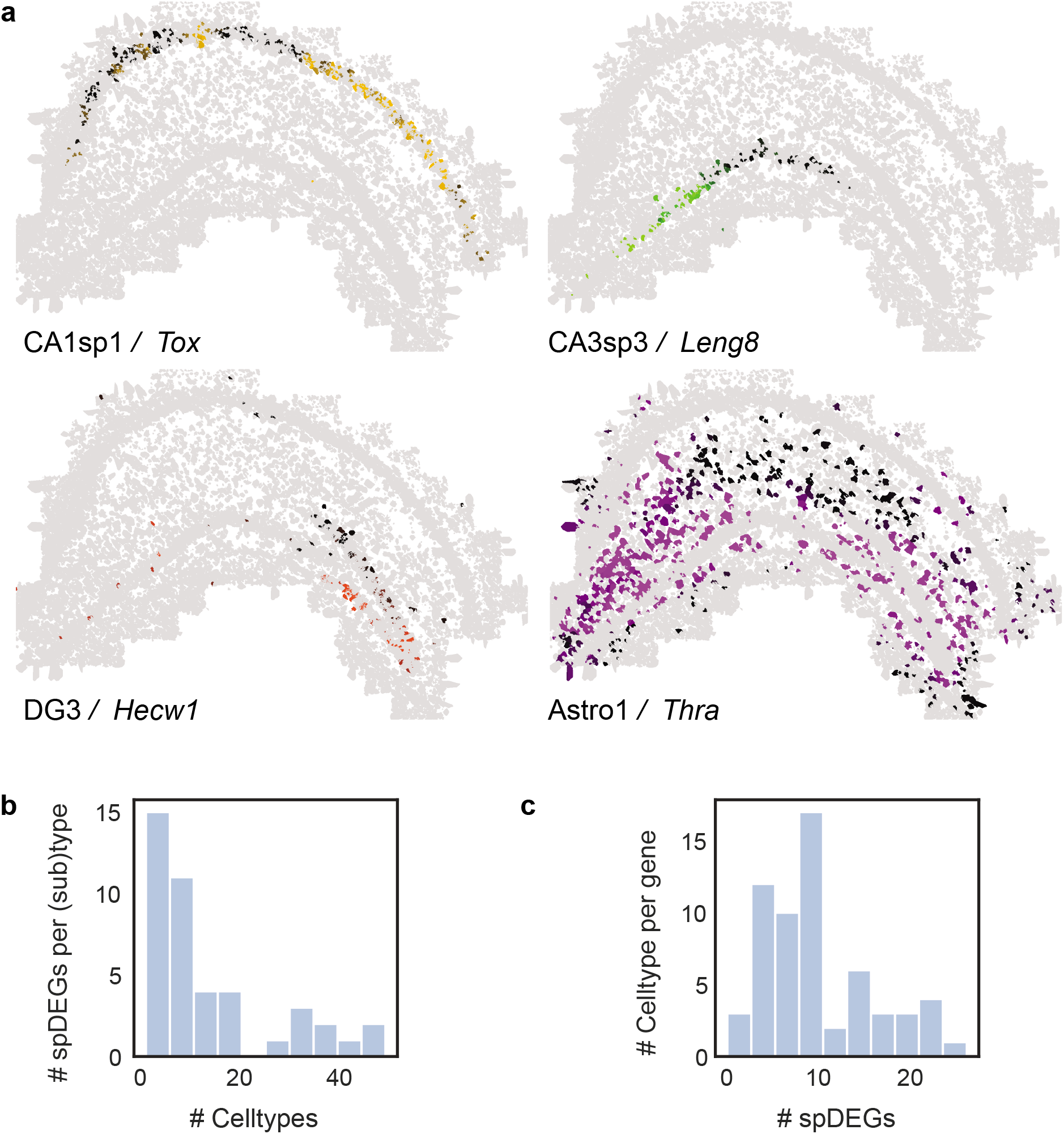
Identification of spatial differential gene expression (spDEGs). **a**. Normalized expression of Tox in CA1sp1, Leng8 in CA3sp3, Hecw1 in DG3, and Thra in Astro1. Scale bar is 500 microns. **b**. Histogram of the number of statistically significant spDEGs (Benjamini-Hochberg corrected FDR < 0.05) in each subtype. **c**. Histogram of the number of subtypes that have an spDEG for each gene.

## Discussion

Spatial transcriptomics provides the coordinates of each transcript without any information on the transcript cell of origin^26^. Here we present JSTA, a new method to convert raw measurements of transcripts and their coordinates into spatial single cell expression maps. The key distinguishing aspect of our approach is its ability to leverage existing scRNAseq based reference cell type taxonomies to simultaneously segment cells, classify cells into (sub)types and assign mRNAs to cells. The unique integration of spatial transcriptomics with existing scRNAseq information to improve the accuracy of image segmentation and enhance the biological applications of spatial transcriptomics, distinguishes our approach from other efforts that regardless of their algorithmic ingenuity are bounded by the available information in the images themselves. As such, JSTA is not a generalist image segmentation algorithm rather a tool specifically designed to convert raw spatial transcriptomics data into single cell level spatial expression maps. We show the benefits of using a dedicated analysis tool through the insights it provides into spatial organization of distinct (sub)types in the mouse hippocampus and the hundreds of newly discovered cell (sub)type specific spDEGs. These insights into the molecular and cellular level structural architecture of the hippocampus demonstrates the types of biological insights provided by highly accurate spatial transcriptomics.

The promise of single cell and spatial biology lends itself to intense focus on technological development and large scale data collection efforts. We anticipate that JSTA will benefit these efforts while at the same time benefit from them. On the technology side, we have demonstrated the performance of JSTA for one specific variant of spatial transcriptomics, MERFISH. However, the algorithm is extendable and could be applied to other spatial transcriptomics approaches that are based on in situ sequencing^27,28,29^, spatial barcoding ^4,30^, and potentially any other spatial “omics” platforms ^31–36^. The benefits of JSTA are evident even with a small number of measured genes. This indicates that it is applicable to a broad range of platforms across all multiplexing capabilities. On the data side, as JSTA leverages external reference data, it will naturally increase in its performance as both the quality and quantity of reference cell type taxonomies improve^37^. We see JSTA as a dynamic analysis tool that could be reapplied multiple times to the same dataset each time external reference data is updated to always provide highest accuracy segmentation, cell (sub)type classification, spDEG identification.

Due to the nascent status of spatial transcriptomics there are many fundamental questions related to the interplay between cell (sub)types and other information gleaned from dissociative technologies and tissue and organ architecture ^38,39^. Our results show that strong co-dependency between spatial position and transcriptional state of a cell in the hippocampus, these results mirror findings from other organs ^40–42^. This codependency supports the usefulness of the reference taxonomies that were developed without the use of spatial information. Agreements between cell type taxonomies developed solely based on scRNAseq and other measurement modalities, i.e. spatial position, corroborates the relevance of the taxonomical definitions created for mouse brain^22^. At the same time, the spatial measurements demonstrate the limitation of scRNAseq. We discovered many spatial expression patterns within most cell (sub)types that prior to these spatial measurements would have been considered biological heterogeneity or even noise but in fact they represent key structural features of brain organization. High accuracy mapping at the molecular and cellular level will allow us to bridge cell biology with organ anatomy and physiology pointing towards a highly promising future for spatial biology.

## Author Contributions

RL, XY, RW developed the algorithm that was implemented by RL. DA and RF designed MERFISH probeset. ZH, RF performed MERFISH measurements and initial image analysis. GZ and FGP performed brain sample preparation.

## Acknowledgment

The work was funded by NIH grant R01NS117148 and T32CA201160

## Source code

https://github.com/wollmanlab/JSTA

## Methods

### Tissue Preparation

B6 mouse was euthanized using carbon dioxide with cervical dislocation. Its brain was harvested and flash frozen in Optimal Cutting Temperature Compound (OCT) using liquid nitrogen. 15 um sections were prepared and placed on pretreated coverslips.

### Coverslip Functionalization

Coverslips were functionalized to improve tissue adhesion and promote gel attachment ^43^. Briefly, 40 mm No.1 coverslips were cleaned with a 50:50 mixture of concentrated 37% hydrochloric acid and methanol under sonication for 30 minutes. Coverslips were silanized to improve gel adhesion with 0.1% triethylamine and 0.2% allyltrichlorosiloxane in methanol under sonication for 30 minutes then rinsed once with chloroform then twice with ethanol. Silanization was cured at 70C for 1 hour. An additional coating of 2% aminopropyltriethoxysilane to improve tissue adhesion was applied in acetone under sonication for 2 minutes then washed twice with water and once with ethanol. Coverslips were dried at 70C for 1 hour then stored in a desiccator for less than 1 month.

### Probe Design and Synthesis

A total of 18 readout probes were used to encode the identity of each gene. Each gene was assigned 4 of the possible 18 probes such that each combination was a minimum hamming distance of 4 away from any other gene. This provides classification that is robust up to 2 bit errors. 80 to 120 encoder probes were designed for each target gene. Encoder probes contained a 30 bp region complementary to the transcript of interest with a melting point of 65C and less than 17 bp homology to off target transcripts including highly expressed ncRNA and rRNA. Probes also contained 3 of 4 readout sequences assigned to each gene. Sequences are available in supplementary material. Probes were designed using modified MATLAB code developed by the Zhuang Lab ^43^. Probes were ordered from custom arrays as a single strand pool. A T7 promoter was primed into each sequence with a limited cycle qPCR to allow amplification through in vitro transcription and reverse transcription^43^.

### Hybridization

Hybridization was performed using a modified MERFISH protocol ^43^. Briefly, Tissue sections were fixed in 4% PFA in 1xPBS for 15 minutes and washed 3 times with 1xPBS for 5 minutes each. Tissue was permeabilized with 1% Triton X-100 in 1xPBS for 30 minutes and washed 3 times with 1x PBS. Tissue was incubated in 30% formamide in 2xTBS at 37C for 10 minutes. Encoding probes were hybridized at 5 nM per probe in 30% Formamide 10% dextran sulfate 1 mg/mL tRNA 1 uM poly T acridite anchor probed and 1% Murine RNAse inhibitor in 2xTBS. A 30 uL drop of this encoding hybridization solution was placed directly on the coverslip and a piece of parafilm was placed on the coverslip to prevent evaporation. Probes were hybridized for 30-40 hours at 37C in a humidity chamber. Tissue was washed twice with 30% formamide in 2xTBS for 30 minutes each at 45C. Tissue was washed 3 times with 2xTBS. Tissue was embedded in a 4% polyacrylamide hydrogel with 0.5uL/mL TEMED 5uL 10% APS and 200nm blue beads for 2 hours (Can expand if wanted). Tissue was cleared with 1% SDS, 0.5% Triton x-100, 1 mM EDTA, 0.8 M guanidine HCl 1% proteinase K in 2xTBS for 48 hours at 37C replacing clearing solution every 24 hours. Sample was washed with 2xTBS and mounted for imaging. Readout hybridization was automated using a custom fluidics system. Sample was rinsed with 2xTBS and buffer exchanged into 10% dextran sulfate in 2xTBS for hybridization. Hybridization was performed in 10% dextran sulfate in 2xTBS with a probe concentration of 3nM per probe. Sample was washed with 10% dextran sulfate then 2xTBS. Sample chamber was filled with a 2mM pca 0.1& rPCO 2mM VRC 2mM Trolox in 2xTBS Imaging Buffer. Sample was imaged at 63X using a custom epifluorescent microscope. After imaging fluorophores were stripped using 50mM TCEP in 2xTBS and the next round of readout probes were hybridized.

### Image Analysis

Image analysis was performed using custom python code^44^. To register multiple rounds of imaging together with sub pixel resolution, fiduciary markers were found and a rigid body transformation was performed. Images were preprocessed using hot pixel correction, background subtraction, chromatic aberration correction, and deconvolution. An 18 bit vector was generated for each pixel where each bit represented a different round and fluorophore. Each bit was normalized so that background approached 0 and spots approached 1. An L2 normalization was applied to the vector and the euclidean distance was calculated to the 18 bit gene barcode vectors. Pixels were classified if their euclidean distance was less than a 2 bit error away from the nearest gene barcode. Individual pixels that were physically connected were merged into a spot. Dim spots and spots that contained 1 pixel were removed.

### Nuclei Segmentation

Nuclei were stained using dapi and imaged after MERFISH acquisition. Each 2D image was segmented using cellpose with a flow threshold of 1 and a cell probability threshold of 0^45^ CITE. 2D masks of at least 10um^2^ area were merged if there was at least 30 percent overlap between frames. 3D masks that were present in less than 5 z frames (2um) were removed.

### Simulation

#### scRNAseq reference preparation

The NCTT was subset to the cells found in the hippocampus, and to the marker genes we had generated in our MERFISH data. Expression levels of simulated genes were taken from scRNAseq reference and were harmonized to qualitatively match the variance observed in measured in MERFISH data. These were then rounded to create a scaled counts matrix. For each of the 133 hippocampal cell types from the NCTT, we computed a mean vector and covariance matrix of gene expression. We additionally computed the cell type proportions in the single cell data for later use in cell type assignment.

#### Creating the cell map

Initially, the cell centers were placed in a 200 X 200 X 30 micron grid, equidistant from one another, with an average distance between cell centers of 4 microns. The cell centers were then moved around in each direction (x, y, z) based on a gaussian function with mean 0 and standard deviation 0.6. Pixels were then assigned to their closest center with a minimum distance of 5 microns and maximum distance of 7 microns. Cell’s with less than 30 pixels were removed due to small unrealistic sizes. To create more realistic and non-round cells, we merged neighboring, touching cells twice. Each cell was assigned a (sub)type uniformly across all 133 types in our dataset. Nuclei were randomly placed within each cell with 20 pixels. Nuclei pixels placed on the border were removed. We simulated 10 independent replicates in each simulation study.

#### Generating cell transcriptional profiles, and placing spots

Each cell’s gene expression profile was drawn from a multivariate gaussian using the mean vector, and covariance matrix computed from the scRNAseq reference. This vector and matrix are cell type specific, and each cell’s gene expression profile is sampled from these cell type specific distributions. The mRNA spots were then placed inside of each cell, slightly centered around the nucleus, but mostly uniform throughout.

#### Simulated data on limited genes

To perform feature selection and extract a limited number of important genes (4, 12, 20, 28, 36, 44), we used a random forest classifier with 100 trees to predict cell types in the reference dataset. The top n important features for classifying cell types were used. Other simulation parameters were the same as above.

### K-Nearest Neighbors (KNN) based Density Estimation Method

We used a KNN approach to estimate density for many genes at each point^46^. The volume required to reach the 5th spot was computed and used to compute the density estimation (Equation 1). Where *r* is the radius to the 5th closest spot of that gene. We repeated this process for all genes.

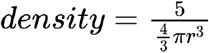

### JSTA Overview

Expectation Maximization (EM) can be used to jointly classify the identity of an observation of interest, while learning the parameters that describe the class distributions. In EM, the object classes are initialized with a best guess. The parameters of the classifying function are learned from this distribution of initialized classes (M-step). The objects are reclassified according to the updated function parameters (E-step). These steps are repeated until the function parameters converge. JSTA is designed with an EM approach for reclassifying border pixels in the 3 dimensional grid of pixels based on their estimated transcriptional densities. First, we initialize the spatial map with watershed, in euclidean space with a maximum radius. Next we classify cell types of the segmented cells based on the computed count matrix. We then randomly sample a fraction of the pixels’ gene expression vectors, and train a pixel classifier (M-step). The pixel classifier is used to reclassify the cell identity of pixels that are at the border between different cell types, or between a cell and empty space (E-step).

### Cell Type Classification

#### Data preparation

To match the distributions of both scRNAseq and MERFISH, we centered and scaled each cell across all genes. We then subsequently centered and scaled each gene across all cells.

#### Cell type classifier

We parameterized the cell type classifier as a neural network, with 3 intermediate layers with 3 times the number of input genes as nodes. We used a tanh activation function with L1 regularization (1e-4) allowing for the influence of negative numbers in the scaled values and parameter space sparsity^47^. Batch normalization was used on each layer^48^, and a softmax activation was used for the output layer^49^ (Table S1).

#### Training the classifier

The network parameters were initialized with Xavier initialization^50^. The neural network was trained with two steps with learning rates of 5e-3 and 5e-4 for 20 epochs each, with batch size of 64, and the Adam optimizer was used ^51^. A 75/25 train validation split was used to tune the L1 regularization parameter and reduce overfitting. We used 75/25 to increase the representation of lower frequency cell classes. Cross entropy loss was used to penalize the model and update parameters accordingly.

### Pixel Classification

#### Pixel classifier

We parameterized the pixel classifier as a neural network with 3 intermediate layers. Each layer was twice the size of the last to increase the modeling power of this network, and indirectly model the other genes not in the MERFISH dataset. Each layer used the tanh activation function and used an l2 regularizer (1e-3). Each layer was centered and scaled with batch normalization, and the output activation was an l2 regularized softmax function (Table S2).

#### Training the classifier

Each time cell types are reclassified, a new network was reinitialized with Xavier initialization. The network was initially trained with learning rates or 1e-3 and 1e-4 for 25 epochs. After the first round of classifying and flipping the assignment of pixels, the network was retrained on a new sample of pixels starting from the previous parameter values. This was then trained with a learning rate of 1e-4 for 15 epochs. All training was performed with the Adam optimizer and a batch size of 64. We used an 80/20 train validation split to help monitor any overfitting that might be occurring, and adjust the hyperparameter selection accordingly. We used cross entropy loss.

#### Identifying border pixels

Border pixels are defined as pixels that are between two cells of different types, or between a cell and empty space. To enhance the smoothness of cells’ borders, we require a border pixel to have 5 of its surroundings be from a different cell, and 2 of its surroundings be from the same cell.

#### Classifying pixels

The trained classifier was then used to estimate the cell type class of border pixels. The pixel classifier outputs a probability vector for each cell type, and the probabilities are scaled by a distance metric based on the distance to the cells’ nuclei that it could flip to. Probabilities less than 0.05 are set to 0. The classification is sampled from that probability vector subset to cell types of its neighbors, and renormalized to 1. If the subset probability vector only contains 0, the pixel identity is set to background. To balance the exploration and exploitation of pixel classification map, we anneal the probability of selecting a non-maximum probability cell type by multiplying the maximum probability by (1 + number of iterations run * *0.05*). If this is selected as 0, complete stochasticity presides, and if it is large, the maximum probability will be selected.

### JSTA Formalization

#### Definitions and background

The gene expression level of *n_c_* cells and *n_p_* pixels are described by the matrices *E_c_* (cells) and *E_p_* (pixels) which are *n_c_*×*m* and *n_p_×m* matrices respectively, where *m* is the number of genes. Likewise, cell type probability distributions of all cells or pixels can be described by matrices. These distributions for cells and pixels are *P_c_* and *P_p_* respectively, represented as *n_c_×k* and *n_p_×k* matrices, where *k* is the number of cell types. We aim to learn *θ* and *ϕ*, such that *f_θ_* and *g_ϕ_*, accurately map from *E_c_* to *P_c_* and *E_p_* to *P_p_*. We used the cross-entropy loss function for penalizing our models.

#### Cell type classification

First, we learn the parameters of *f_θ_* by:

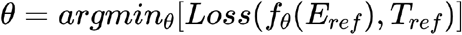

Where *E_ref_* is an *n_ref_×m* gene expression matrix representing the harmonized NCTT data and *T_ref_* is an *n_ref_* vector of cell types labels provided by NCTT. We then use the newly learned mapping to infer the cell type probability distributions in the initialized dataset *E_c_* with:

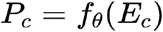

We classify each cell as the highest classification probability for that cell:

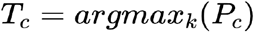

Where *T_c_* are the predicted cell types for each of the cells in the matrix *E_c_*.

#### Joint pixel and parameter updates

We initialize the labels *T_p_* for all pixels based on the current segmentation map that assigns pixels to cells. We then learn the parameters of the mapping function *g_ϕ_* (maximization). Learning is performed by updating the parameters of the mapping function *g_ϕ_* with:

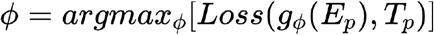

The updated mapping function is then used to infer the probability of observing a type *T_p_* given expression *E_p_* in all pixels:

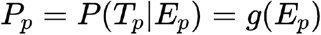

The next step is to update *P_p_* based on spatial proximity to cells of each type. Using the notation *q* for the vector of probabilities of a single pixel (*q* = *P_pj_* = [*q*_0_ …, *q_i_*, …, *q_k_*]) we next update the elements in the vector q based on neighborhood information. We scaled the values of *q_i_* based on its distance from the nuclei and its neighbors. *q′* is intermediate in the calculation that does not represent true probabilities.

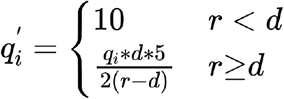

Where *r* is the distance from the nucleus of the closest cell of cell type *i*, *d* is the distance threshold for which a pixel should automatically be assigned to that nucleus. The values 10 and 5 were determined empirically to modify the sharpness of probability decline based on distance. 10 was chosen to be much bigger than probabilities produced by *g_ϕ_* and 5 was chosen to allow the probability to decay to half over 5*d*.

We then only kept probabilities for cell types of neighboring cells:

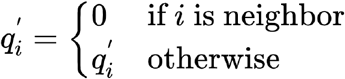

We then used the intermediate q’ to recalculate the pixel type probabilities:

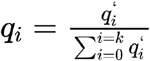

The updated values per cell (*q_j_*) are then used to update the probability matrix *P_p_*. The type per pixel (*T_p_*). The assignment of pixel to cells is then stochastically assigned according to: the inferred probability *P_p_* per pixel basis.

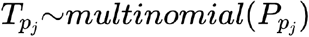

We then repeat updating *g_ϕ_* and *T_p_* until convergence.

### Segmentation

#### Density estimation

The 3 dimensional space was broken into a grid of pixels with the edge of each pixel 2 microns in length (1 micron in simulation). The density was estimated at the center of each pixel, for each gene. The volume required to reach 5 mRNA molecules was used as the denominator of the density estimation.

#### Segmentation with JSTA

The cell assignment map was initialized with watershed on the distance transform with a maximum distance from the nucleus of 2 microns. The cells were only classified once. The pixel classifier was trained 6 times (5 in simulation) on 10% of the pixels excluding pixels without assignment. After each training step, we reassigned pixels for 10 iterations (5 in simulation). The lowest probability kept in the predicted pixel assignment vector was 0.05 (0.01 in simulation.

#### Segmentation with watershed

The overall gene density was the sum of each gene in a given pixel. To smooth the range of the density, we log_2_ transformed the density values. Log transformed density values less than 1 were masked. The segmentation used the nuclei locations as seeds and watershed from the skimage python package, with *compactness* of 10. Using compactness of 10 was the highest performing value for watershed. A watershed line was used to separate cells from one another.

### Evaluation of Segmentation in Simulated data

mRNA spot call accuracy was evaluated at different taxonomic levels. For a given cell the accuracy was defined as the number of mRNA spots correctly assigned to that cell divided by the total number of mRNA spots assigned to that cell. To match the algorithm’s ability to segment based on cell type information, RNAs that were assigned to a neighboring cell of the same (sub)type were also considered correct assignment. The overall segmentation accuracy was the mean accuracy across all cells in a given sample. To evaluate accuracy at different levels, we utilized the NCTT dendrogram. We used dendrogram heights at 0 through 0.8 with a step size of 0.05 (133, 71, 32, 16, 11, 8, 5, 4, 3, 2 cell types).

### Correlation of Segmented MERFISH with scRNAseq

The NCTT scRNAseq data was subset to the marker genes in our MERFISH dataset. Cells in the segmented MERFISH dataset were assigned to canonical hippocampus cell types (Astrocyte, CA1 pyramidal neuron, CA2 Pyramidal neuron, CA3 Pyramidal, Dentate Gyrus, Inferior temporal cortex, Macrophage, Oligodendrocyte, Subiculum, Interneuron) based on their high resolution cell type classification. In each cell type the average expression in each gene was calculated. Only genes were kept that had an average expression of at least 5 counts in one of the cell types. Values were centered and scaled across all cell types. The Pearson correlation was computed for each gene for the matching cell types between scRNAseq and MERFISH.

### Distribution of High Resolution Celltypes in the Hippocampus

CA1 and CA3 subtypes were projected onto the lateral medial axis. The smoothed density across this dimension was plotted for each of the subtypes.

### Colocalization of High Resolution Celltypes

Significant colocalization of subtypes was determined through a permutation test. First, the 20 nearest cell types around each cell were determined. We counted the number of cells from each type that surround each cell type and computed the fraction of neighbors coming from each subtype. This created a matrix with the fraction of colocalizations per cell between each cell type combination. We then permuted the labels of the cell types 1000 times, and recomputed this interaction matrix to create a null distribution. For each cell type colocalization, we determined the percentage of colocalizations in the null distribution that are higher than the true colocalization number to create a p-value for each colocalization. We corrected for multiple testing with the benjamini hochberg procedure and determined significance using FDR < 0.05.

### Identification of Spatial Differential Gene Expression (spDEGs)

spDEGs were calculated in cell types with more than 40 cells. Within each cell type, we computed a local expression of each non-marker gene for each cell. The local expression was the mean expression of a gene in the cell and its 9 nearest neighbors. We then built a null distribution by permuting gene expression values within the cell type, and repeating the local expression process for 100 permutations. Determining if a gene was spatially differentially expressed, we compared the null distribution within a cell type with the true distribution of local expression using a chi squared test. We corrected for multiple testing with benjamini hochberg procedure and determined significance using FDR < 0.05.

### Python packages used

python (3.8.3), numpy (1.18.5), pandas (1.0.5), matplotlib (3.2.2), scipy (1.5.0), scikit-learn (0.23.1), scikit-image (0.16.2), tensorflow (2.2.0). seaborn (0.10.1)

## Supplementary Figures

**Figure S1.**
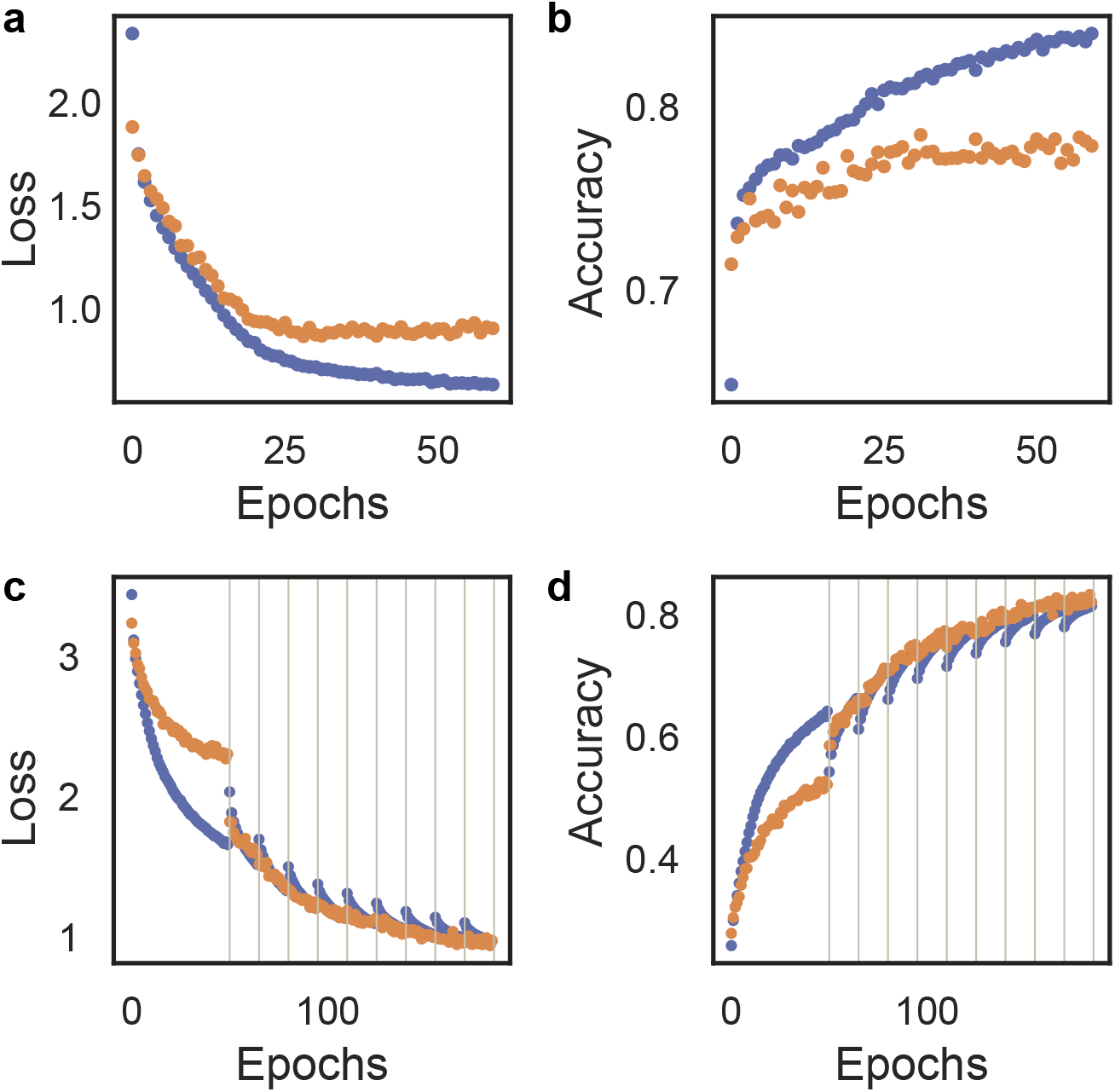
Loss and accuracy of cell type (**a-b**) and pixel (**c-d**) classifier during training for the train (blue) and validation (orange) data sets. **a-b**. Cross entropy (**a**) loss and accuracy (**b**) during training cell type classifier. **c-d**. Cross entropy loss (**c**) and Accuracy (**d**) during training of the pixel classifier. Black lines indicate new training iteration after pixel reassignment.

**Figure S2.**
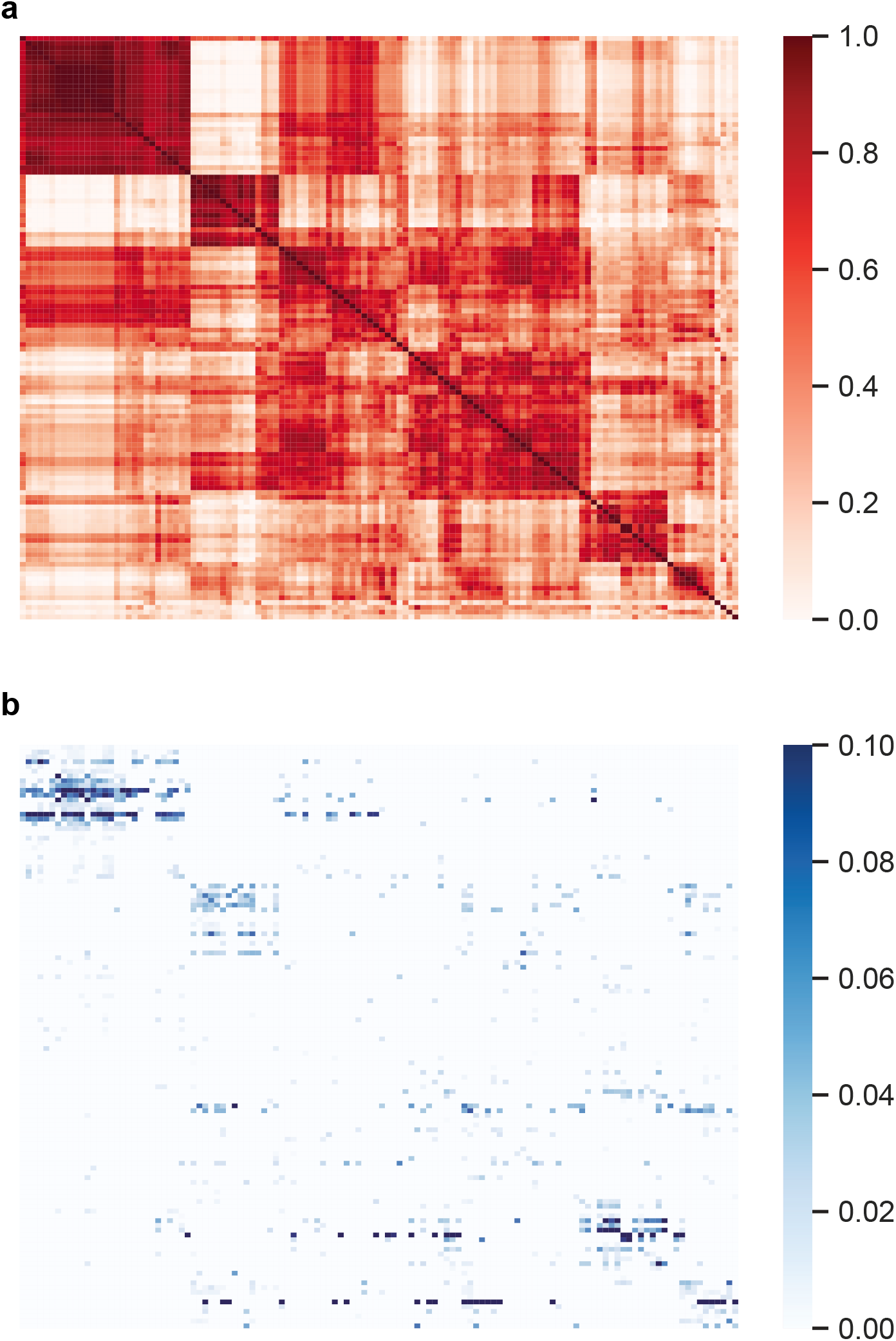
Correlation structure of cell types compared to their colocalization. Cell types with more than 10 cells were included. **a**. pearson correlation of 122 (sub)types across 83 marker genes. **b**. Frequency of neighbors between each of 122 (sub)types. Only significant (FDR < 0.05) colocalizations are shown. Labels and values are detailed in supplementary table 3 and 4.

**Figure S3.**
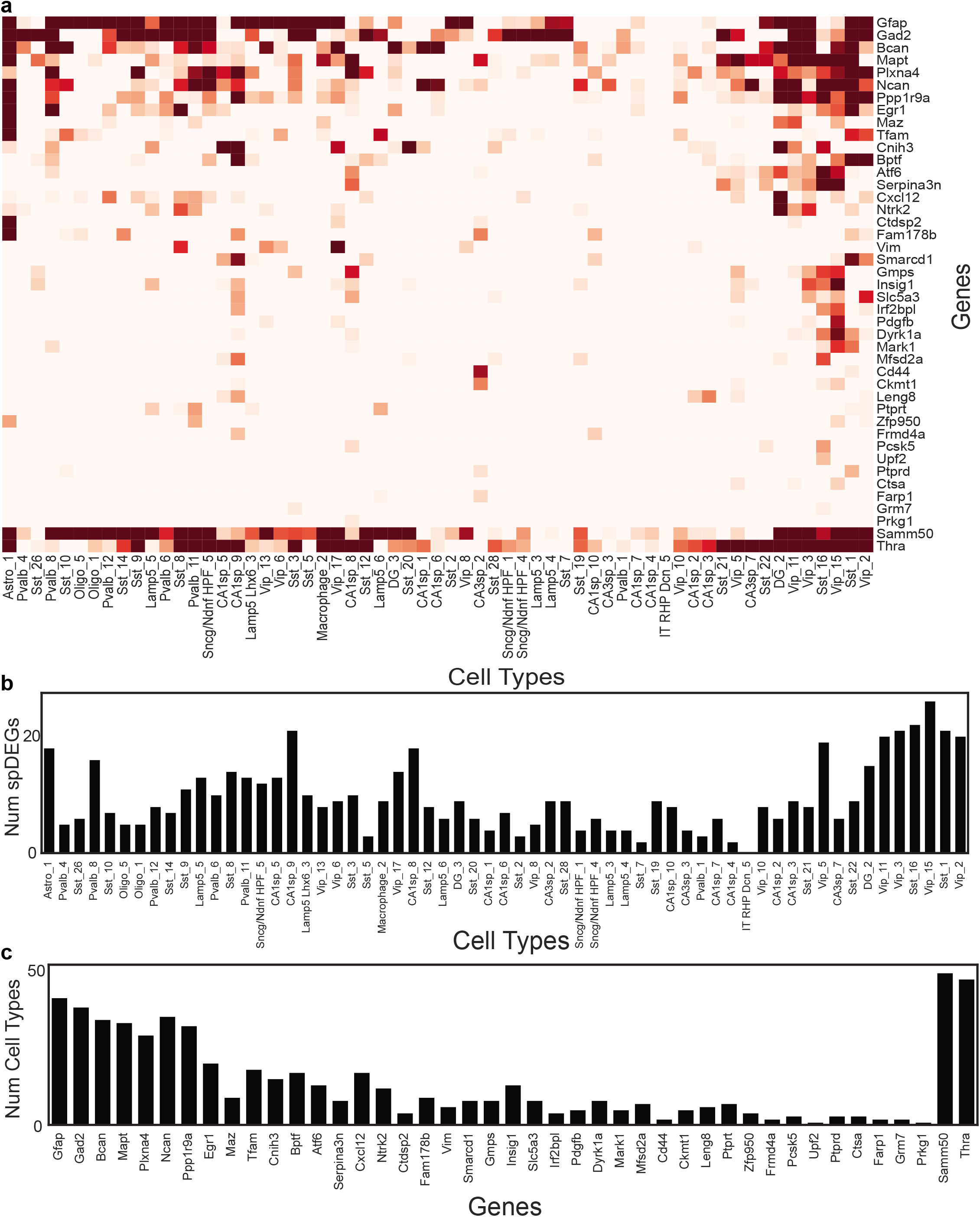
Identification of spatial differentially expressed genes (spDEGs). **a**. 43 genes across 61 cell types show significant spDEGs. Heatmap values correspond to −log_2_(p-value). **b**. Number of spDEGs in each of the 61 cell types. **c**. Number of cell types with each of the 43 spDEGs.

## Supplementary Tables

**Table S1.** Cell type classifier architecture. The network was initialized with Xavier initialization. Learning rates of 5e^−3^ and 5e^−4^ for 20 epochs each. A batch size of 64 was used. The Adam optimizer was used to update parameters. Cross entropy loss was used.

| Layer Type | Num Nodes | Activation | Regularization |
| --- | --- | --- | --- |
| Input | 83 | - | - |
| Dense | 249 | tanh | L1 ($5e^{-3}$) |
| Batch Normalization | - | - | - |
| Dense | 249 | tanh | L1 ($5e^{-3}$) |
| Batch Normalization | - | - | - |
| Output | 133 | softmax | L1 ($5e^{-3}$) |

**Table S2.** Pixel classifier architecture. The network was initialized with Xavier initialization. Initially the model was trained for 25 epochs with 1e^−4^ and 1e^−3^ learning rate. Subsequent updates were done on 15 epochs with a learning rate of 1e^−4^. We used the Adam optimizer to update parameters. Cross entropy loss was used.

| Layer Type | Num Nodes | Activation | Regularization |
| --- | --- | --- | --- |
| Input | 83 | - | - |
| Dense | 166 | tanh | L1 ($1e^{-3}$) |
| Batch Normalization | - | - | - |
| Dense | 166 | tanh | L1 ($1e^{-3}$) |
| Batch Normalization | - | - | - |
| Dense | 332 | tanh | L1 ($1e^{-3}$) |
| Batch Normalization | - | - | - |
| Output | 133 | softmax | L1 ($1e^{-3}$) |

